# MAGIC: Methylation Analysis with Genomic Inferred Contexts

**DOI:** 10.1101/2025.11.19.689248

**Authors:** Jiaqi Han, Michael Thompson, Matteo Pellegrini

## Abstract

**Motivation:** Whole-genome bisulfite sequencing (WGBS) enables base-resolution methylation profiling but poses statistical challenges for differential methylation analysis. While existing methods have made important progress using uniform dispersion shrinkage across CpGs, there remains opportunity to better capture methylation heterogeneity across the genome. There is a need to account for context-specific methylation variability and to improve detection accuracy under realistic sequencing constraints.

**Results:** We propose MAGIC (Methylation Analysis with Genomic Inferred Contexts), a beta-binomial mixture model that learns genomic contexts directly from methylation data. MAGIC models genomic heterogeneity through context-specific dispersion shrinkage and detects differential methylation via two complementary tests, including a Wald test and Bayes factor test. Simulations across various coverage and effect sizes demonstrate that MAGIC improves sensitivity and false-positive rate compared with DSS, providing a robust framework for differential methylation analysis in WGBS data.

**Availability:** The source code of MAGIC is available at https://github.com/Pickledzebra/MAGIC.

**Supplementary information:** Supplementary data are available at bioRxiv.

## 1 Introduction

DNA methylation is an important epigenetic modification that regulates gene expression and contributes to the maintenance of genomic stability (Bird *et al*. 2002). It plays essential roles in genome imprinting, cellular differentiation, and aging (Li *et al*. 1993; Jones *et al*. 2012; Seale *et al*. 2022). Aberrant methylation patterns are closely associated with various human diseases, including cancer, pregnancy disorders, and neurological abnormalities (Gimeno *et al*. 2025; De Borre *et al*. 2023; Moore *et al*. 2025).

Whole-genome bisulfite sequencing (WGBS) provides single-base resolution measurements across the entire genome. Following sodium bisulfite treatment, unmethylated cytosines are converted into uracils and subsequently read as thymines, whereas methylated cytosines remain unchanged (Lister *et al*. 2009). The development of these high-throughput sequencing technologies has enabled differential methylation analysis, including the identification of differentially methylated loci (DML) and differentially methylated regions (DMR). However, WGBS data often suffer from low sequencing depth and limited biological replicates due to the high experimental cost, and methylation counts exhibit variation arising from both biological and technical noise (Ziller *et al*. 2015; Sun *et al*. 2014). These characteristics pose significant statistical challenges for differential methylation analysis.

Identifying DML between conditions represents a central objective in methylation analysis, as these loci reveal crucial epigenetic changes associated with disease, developmental process, and aging (Shafi *et al*. 2018). Several computational methods have been developed to detect DML. The simplest strategy involves applying statistical tests like Fisher’s exact test and t-test between groups without considering biological variability. More advanced approaches, such as MethylSig (Park *et al*. 2014) and DSS (Park *et al*. 2016), treat the count data as a beta-binomial distribution, considering both biological and technical noise. However, their performance often declines when applied to low coverage datasets. Alternatively, methods like BSmooth (Hansen *et al*. 2012) and BiSeq (Hebestreit *et al*. 2013) use smoothing or aggregation methods to enhance detection sensitivity by borrowing information from neighboring CpG sites. While these methods pool information to stabilize dispersion estimates, they assume a uniform distribution of methylation dispersion, which may not accurately reflect the true biological diversity (Piao *et al*. 2021). Another category of methods, including MethylKit (Akalin *et al*. 2012), RADMeth (Dolzhenko *et al*. 2014) and DMAP (Stockwell *et al*. 2014), adopts regression frameworks. The last category of tools utilizes Hidden-Markov Models (HMM), like HMM-Fisher (Sun *et al*. 2016), to model methylation states and their transitions along the genome. These methods can handle low coverage data more effectively than binomial-based approaches, but are computationally intensive and often lack generalizability. In summary, although a wide range of tools have been developed, accurately detecting DML under low-coverage conditions remains a challenge.

One major limitation is that under low coverage, the dispersion of methylation levels at CpG sites is often overestimated. Among existing methods, DSS has demonstrated relatively strong performance in detecting differential methylation through global dispersion shrinkage (Piao *et al*. 2021). However, promoters, gene bodies, and intergenic regions exhibit distinct methylation patterns (Jones *et al*. 2012). CpG-dense regions such as CpG islands generally exhibit low methylation with minimal variation, whereas CpG shores display more variable methylation patterns (Schultz *et al*. 2015). Therefore, incorporating local heterogeneity in methylation dispersion across diverse genomic contexts may further improve the detection performance. This opportunity highlights the need for context-specific dispersion shrinkage.

Mixture models are particularly effective for modeling complex data where observations arise from a combination of distinct underlying distributions (McLachlan and Peel, 2000). Bayesian inference further enhances this framework by quantifying uncertainty through posterior inference and latent assignments (Gelman *et al*. 2013). Unlike frequentist methods, which typically provide point estimates and rely on large-sample approximations, Bayesian inference produces full posterior distributions and allows hypothesis comparison through Bayes factors. Because it does not depend strictly on asymptotic theory, Bayesian inference can be useful when data are sparse or sample sizes are small (Kruschke *et al*. 2015).

In the context of DNA methylation analysis, Bayesian mixture models have been widely applied to address the heterogeneity inherent in methylation data. For example, scMET (Kapourani *et al*. 2021) demonstrated that Bayesian mixture models can effectively identify subpopulations of cells in single-cell Bisulfite Sequencing (scBS-seq) data. BayesDiff (Gu *et al*. 2022) applied a nonparametric Bayesian approach to model complex correlation structures in EPIC array data, enhancing differential methylation detection across cancer cohorts. BMM provided an effective strategy for differential methylation analysis, offering biologically interpretable methylation states without the need for data transformation (Majumdar *et al*. 2022). However, these models have not yet been adapted for detecting differential methylation in WGBS data, which present distinct statistical challenges.

Here, we propose a novel Bayesian mixture model, MAGIC (Mixture-based Analysis with Genomic Inferred Contexts), which integrates local methylation features to model heterogeneity in dispersion. Extending the dispersion shrinkage framework established by DSS, our model infers contexts directly from observed methylation dispersion across samples rather than relying on predefined genomic annotations. By probabilistically assigning CpG sites to components and conducting component-specific dispersion shrinkage, MAGIC provides more accurate estimation of methylation dispersion and improves the detection of differential methylation sites under low coverage and low effect size conditions. With synthetic datasets, we demonstrate that MAGIC achieves superior false positive control and detection performance compared with existing methods.

## 2 Methods

### 2.1 Model Architecture

Whole-genome bisulfite sequencing (WGBS) provides base-resolution methylation measurements, but the data are often sparse and unevenly covered across CpG sites, posing challenges for accurate estimation of methylation variability. Conventional models like DSS lay the foundation for dispersion shrinkage using a global dispersion parameter. MAGIC extends this framework by introducing a finite beta-binomial mixture model in which each mixture component represents a genomic context characterized by distinct methylation mean and dispersion. CpG sites are probabilistically assigned to these components, enabling context-specific dispersion shrinkage and flexible modeling of multimodal methylation distributions across the genome.

For CpG site *i* in sample *j*, let *m*_*ij*_ denote the number of methylated reads and *n*_*ij*_ the total coverage. The observed methylation proportion is:

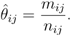

To account for cellular heterogeneity, the true methylation level *θ*_*ij*_ is modeled as a Beta distribution:

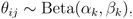

and given this level, the observed methylated counts follow a Binomial distribution:

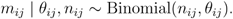

Integrating out *θ*_*ij*_ gives a Beta-Binomial likelihood:

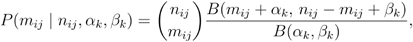

where B(·, ·) is the beta function. The mean and dispersion of the Beta-Binomial distribution can be calculated as:

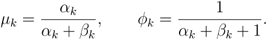

Where *μ* represents the expected methylation proportion and *ϕ* controls overdispersion relative to the Binomial variance (Griffiths, 1973).

The variance can be decomposed as:

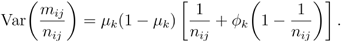

MAGIC assumes that each CpG site is probabilistically associated with the components, each characterized by distinct Beta distributions *B(α*_*k*_, *β*_*k*_) and mixing proportion *π*_*k*_. The optimal number of components k is determined by training models with different k values on a training subset and selecting the k with the lowest BIC on a held-out test set. This leads naturally to a finite mixture model:

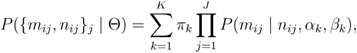

Where Θ = { *π*_1,_ … *π*_*K*,_ *α*_1_, *β*_1,_ …, *α*_*K*_, *β*_*K*_ **}**.

This mixture structure allows MAGIC to learn methylation contexts directly from data without relying on predefined genomic annotations, enabling adaptive, context-specific shrinkage of dispersion and improved modeling of heterogeneous methylation landscapes.

### 2.2 Parameter Estimation

Model parameters were estimated by maximizing the observed data log-likelihood *l*:

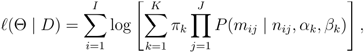

subject to *α*_*k*_, *β*_*k*_ > 0, and 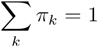

To ensure numerical stability and enforce parameter constraints, optimization was performed in unconstrained space using the transformations:

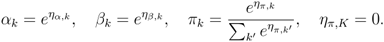

Gradients were computed analytically. The posterior responsibility *r*_*ik*_ of site *i* for component *k* is:

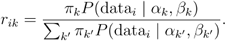

The gradient of the log-likelihood with respect to *α*_*k*_ is:

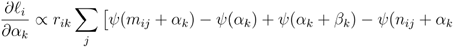

where *ψ(*⋅) is the digamma function. All derivatives were transformed via the chain rule to the unconstrained parameterization.

Optimization was conducted using the L-BFGS-B quasi-Newton algorithm (Byrd *et al*. 1995) with multiple random initializations. The best run was selected based on the maximum log-likelihood, and components were ordered by increasing mean methylation *μ*_*k*_ for interpretability.

Model selection for the optimal number of components *k* was guided by the Bayesian Information Criterion (BIC) (Schwarz *et al*. 1978):

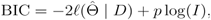

where *p* is the number of free parameters and *I* is the number of CpG sites. AIC and held-out log-likelihood were used as secondary validation criteria (Akaike *et al*. 1974).

### 2.3 Posterior Inference

After parameter estimation, MAGIC computes the posterior probability that CpG site *i* belongs to each component *k*:

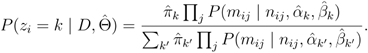

The soft assignments *P* (z_*i*_ = *k*) provide probabilistic context memberships, while hard assignments:

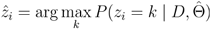

yield discrete context labels for downstream analysis.

Each inferred context *k* is characterized by mean methylation 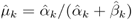, dispersion 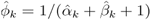, and mixing proportion 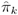.

The mixture model decomposition of methylation patterns provides a biologically interpretable view of epigenomic regulation. Unmethylated contexts (*μ* ≈ 0.05) correspond to active promoters, facilitating gene transcription, whereas lowly methylated regions (*μ* ≈ 0.3) mark enhancers and regulatory elements driving cell-type specific expression. Intermediately methylated contexts (*μ* ≈ 0.5) often reflect dynamic chromatin states, including bivalent domains or allele-specific methylation. Gene bodies with moderate-high methylation (*μ* ≈ 0.7) are associated with transcriptional elongation, and highly methylated regions (*μ* > 0.9) correspond to repressed repetitive elements and heterochromatin.

The estimated dispersion (*φ*_*k*_) indicates regulatory flexibility, with high-dispersion contexts reflecting dynamic epigenetic modulation and low-dispersion contexts representing stable maintenance. The mixing proportions (*π*_*k*_) quantify the relative contribution of each methylation context, allowing assessment of global regulatory shifts across developmental stages, disease states, and perturbations.

### 2.4 Statistical Testing

To identify differential methylation between experimental groups, MAGIC implements two complementary statistical frameworks, including a frequentist Wald test and a Bayes factor test. Both approaches leverage posterior-weighted methylation estimates derived from the fitted mixture model, thereby integrating uncertainty across genomic contexts.

MAGIC implements a Wald test that integrates uncertainty across components. For each site, the Wald statistic is calculated as the estimated methylation difference divided by its standard error.

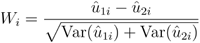

Under the null hypothesis of no methylation difference, this statistic approximately follows a standard normal distribution. Two-sided p-values are obtained from the normal cumulative distribution.

In addition to the Wald test, MAGIC implements a Bayesian test based on the Dirichlet-multinomial mixture model. For each site, let *n*_*pooled*_, *n*_*1*_, and *n*_*2*_ denote the posterior-weighted counts of reads across the *k* mixture components in the pooled, group 1, and group 2 samples, respectively. With component-specific pseudo-counts *α*_*0*_, the marginal likelihoods under the null (*M*_*0*_) and alternative (*M*_*1*_) models are:

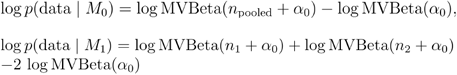

where MVBeta(⋅) denotes the multivariate Beta function. The Bayes factor comparing *M*_*1*_ to *M*_*0*_ is then:

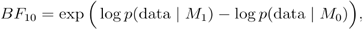

and the posterior probability that the site is differentially methylated is computed as:

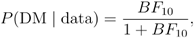

which corresponds to a prior probability *π*_*a*_ = 0.5. This approach naturally propagates uncertainty from the mixture components into inference, and ensures that both the Bayes factor and posterior probability reflect the full mixture-model uncertainty.

To control false discoveries, MAGIC applies the Benjamini–Hochberg procedure (Benjamini *et al*. 1995) to adjust p-values from the Wald or Bayes factor tests. For each site, the results table reports the posterior responsibilities for all components, the posterior-weighted methylation estimates for control and treatment groups, the estimated effect size, the p value of the Wald test, the Bayes factor, and the FDR-adjusted significance level.

All computation in MAGIC is performed using responsibility-weighted aggregation, ensuring that uncertainty from component assignments is propagated into inference. The variance approximations rely on large-sample assumptions. By default, the Wald test serves as the frequentist test, while the Bayes factor offers a complementary Bayesian evidence measure. Together, these two approaches allow MAGIC to balance statistical power and false discovery control under realistic bisulfite sequencing conditions.

### 2.5 Computational Efficiency

MAGIC is optimized for genome-scale analysis through OpenMP multi-threading for parallelization and chunked processing for memory efficiency. All likelihood calculations are performed in log-space with specialized mathematical functions to ensure numerical stability. To address the multi-modal likelihood surface, MAGIC performs multiple optimization runs with different random initializations and selects the solution with maximum log-likelihood. For a typical analysis of approximately 100,000 CpG sites using 4 CPU cores, model optimization with K=3 components requires 2-5 minutes, and subsequent differential testing completes within 5-10 minutes. The computational time scales approximately linearly with the number of CpG sites and mixture components, making MAGIC suitable for whole-genome bisulfite sequencing analysis.

### 2.6 Data Simulation

To systematically evaluate the performance of MAGIC, we developed a method to generate synthetic WGBS datasets that replicates the key characteristics of real bisulfite sequencing data while embedding true DMLs. This method enables comprehensive evaluation of model performance under realistic sequencing settings.

We modeled genome-wide methylation heterogeneity using a beta-binomial mixture model with 3 and 5 components. Each component followed a distinct Beta distribution, Beta(10, 90), Beta(10, 10), and Beta(90, 10), corresponding to hypo-, intermediate-, and hyper-methylated states with mean methylation levels of 0.1, 0.5, and 0.9, respectively. A symmetric dispersion pattern was applied, with lower dispersion at extreme methylation states and higher dispersion at intermediate levels, consistent with biological observations (Rakyan *et al*. 2011). CpG sites were randomly assigned to components with uniform mixing proportions *π*_*k*_ = *1/K*.

Sequencing coverage was drawn from a negative binomial distribution, Coverage_ij_ ∼ NB(*μ, φ*), with mean coverage *μ* and dispersion *φ*. For each CpG site *i* in sample *j*, true methylation proportions *p*_*ij*_ were sampled from the site’s assigned beta distribution, and observed methylated counts were generated according to M_ij_ ∼ Binomial(Coverage_ij_, *p*_*ij*_). This two-stage process captures both biological and technical variation (Sun *et al*. 2014).

Samples were equally divided into control and treatment groups. A random subset of CpG sites was defined as true DMLs, and methylation levels in the treatment group were adjusted by an effect size (delta), *p*_*ij*_ ^*treatment*^ *= p*_*ij*_ ^*control*^ *+ delta*. Sites with extreme baseline methylation (*p*_*ij*_ ^*control*^ > 0.8 or *p*_*ij*_ ^*control*^ < 0.2) were excluded from shifts in the corresponding direction to avoid boundary effects. Methylated counts for treatment samples were regenerated using the shifted proportions. Component-wise methylation distributions are shown in **Supplementary Figure S1-S2**.

We generated datasets containing 100,000 CpG sites distributed across human chromosomes, with 20% of these sites representing true DMLs. Each combination of parameters (*k*, delta, coverage) constituted one simulation scenario, with 10 independent replicates generated per scenario using different random seeds. Ground truth DML annotations were recorded for performance evaluation, and synthetic data were stored as BSseq objects (Hansen *et al*. 2012). All simulations were initialized with fixed random seeds to ensure reproducibility.

### 2.7 Model Evaluation

We evaluated MAGIC’s performance by comparing it against DSS and a baseline approach. The baseline method ranked CpG sites by the absolute difference in mean methylation between groups, serving as a non-statistical benchmark. For MAGIC, the Bayes factor test and the Wald test were used to identify differential methylation.

False positive control was assessed by generating null simulation datasets with no true DMLs. For each method, p-values from the Wald test were obtained for all CpG sites, and quantile-quantile (QQ) plots were constructed to compare the observed p-value distribution with the expected uniform distribution under the null hypothesis.

We systematically assessed model performance across a grid of simulations, varying the number of mixture components (*k* ∈ {3, 5}), effect size (delta ∈ {0.05, 0.1, 0.2}), and coverage (*μ* ∈ {5×, 10×, 20×}). Detection performance was assessed using receiver operating characteristic (ROC) curves, which plot the true positive rate against the false positive rate across varying decision thresholds. True DML status was determined from ground-truth annotations recorded during data generation. For DSS and MAGIC’s Wald test, CpG sites were ranked by descending Wald test statistics. For the MAGIC Bayes factor test, sites were ranked by normalized Bayes factors (Wakefield *et al*. 2008). The baseline method ranked sites by the absolute mean methylation difference between groups. The area under the curve (AUC) was computed for each method in each simulated dataset. All statistical analyses and figure generation were performed in R. ROC curves were generated using the pROC package (v1.19.0.1, Robin *et al*. 2011), and data visualization was conducted using ggplot2 (v3.5.2, Wickham *et al*. 2016).

## 3 Results

### 3.1 Overview

The overall workflow of MAGIC is illustrated in **Fig. 1**. The workflow begins with preprocessed WGBS data, such as output from BSBolt (Farrell *et al*. 2021), converted to BSseq objects containing methylation and read count matrices (**Fig. 1A**). MAGIC then applies a beta-binomial mixture model and decomposes genome-wide methylation heterogeneity into distinct components without requiring prior genomic annotation (**Fig. 1B**). Each component is characterized by its mean methylation level and dispersion parameter, which reflects biological variability beyond technical noise. These inferred contexts correspond to functionally distinct genomic regions, such as unmethylated CpG islands, variably methylated enhancers, and stably methylated heterochromatin (**Fig. 1C**). MAGIC performs context-specific dispersion shrinkage to improve parameter estimates and implements two complementary differential methylation tests, including Bayes factor test and Wald test (**Fig. 1D**). To benchmark model performance, we generated synthetic datasets with known ground truth (**Fig. 1E**) and evaluated detection performance using ROC plots (**Fig. 1F**).

**Fig. 1.**
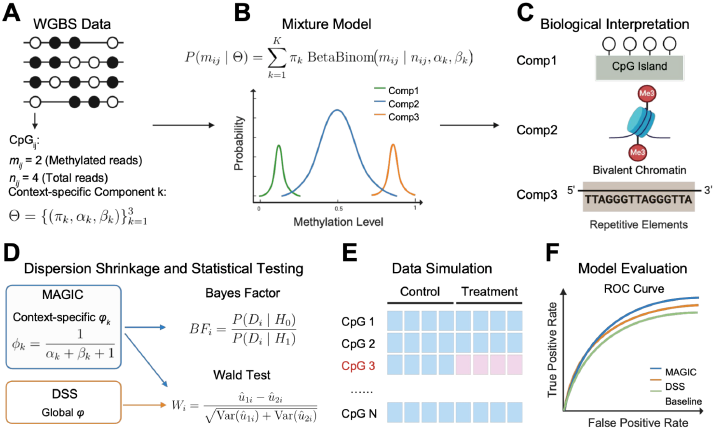
Schematic Overview of the MAGIC Model. (A) Illustration of WGBS data distribution. The count data at each CpG site are modeled using a beta-binomial distribution, where m denotes the number of methylated reads and *n* represents the total coverage. (B) Beta-binomial mixture model. The overall data are represented as a mixture of components, each characterized by distinct weights and beta distribution parameters (π, α, β), resulting in different methylation levels and dispersions. CpG sites are assigned to components according to their dispersion. (C) Biological interpretation of mixture components with varying dispersion levels. (D) Comparison between MAGIC and DSS in dispersion shrinkage (left) and statistical testing (right). MAGIC applies context-specific dispersion shrinkage and provides both Bayes factor and Wald tests for differential methylation detection. (E) Simulation design. 20% of CpG sites were randomly assigned as true DMLs. (F) Performance evaluation using ROC plots. *Created with BioRender*.*com*.

### 3.2 Null Simulation

To evaluate the calibration of MAGIC under the null hypothesis, we simulated datasets without any true differential methylation between two groups, using 10× coverage and 5 biological replicates per group. **Fig. 2** presents a Q–Q plot comparing the observed and expected –log p-values. Deviations above the diagonal line indicate inflation of Type I error, while close alignment suggests proper calibration consistent with the expected uniform distribution.

**Fig. 2.**
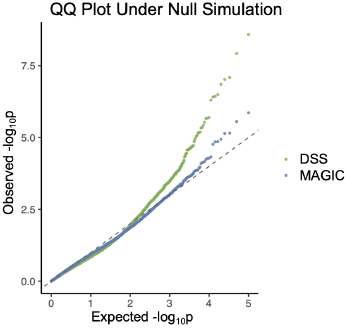
MAGIC produced well-calibrated results under null simulations. MAGIC showed minimal deviation from expected diagonal line (blue), while DSS produced more false positives (green). The dataset was generated with delta = 0, 3 components, 10x coverage, and 5 samples per group.

As shown in **Fig. 2**, MAGIC Wald test produced well-calibrated p-values that closely followed the diagonal line. The observed distribution showed minimal deviation from the theoretical expectation, with no evidence of systematic inflation even in the extreme tails of the distribution, where false positive control is most challenging. In contrast, DSS exhibited an upward deviation, particularly in the upper tail of the distribution. These results demonstrate that MAGIC maintains appropriate statistical calibration under null conditions, reflecting its robustness in avoiding spurious discoveries when no true signal exists.

### 3.3 Performance comparisons on simulated datasets

We then assessed the performance of MAGIC in comparison to DSS and the baseline delta-threshold approach across multiple simulation scenarios. **Fig. 3** shows representative ROC curves obtained with five samples per group and the ROC plots for all simulation settings are provided in **Supplementary Fig. 3–4**. ROC curves displayed the true positive rate (sensitivity) against the false positive rate (1 - specificity) across all possible decision thresholds, with curves closer to the upper-left corner indicating superior performance. The AUC value provides a summary metric ranging from 0.5 (random performance) to 1.0 (perfect discrimination), with higher values indicating better overall ability to distinguish true differential methylation from the noise (Fawcett *et al*. 2006).

**Fig. 3.**
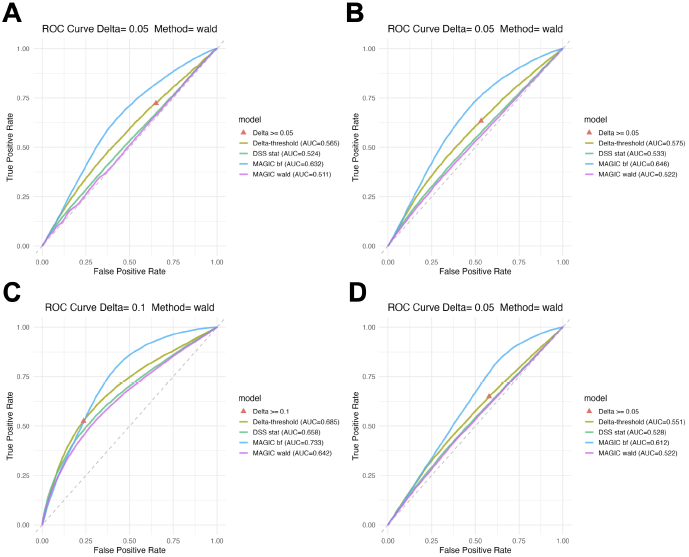
Comparison of model performance on simulated datasets. (A) delta = 0.05, 3 components, 5× coverage. (B) delta = 0.05, 3 components, 10× coverage. (C) delta = 0.1, 3 components, 10× coverage. (D) delta = 0.05, 5 components, 10× coverage. All datasets included 5 samples per group. The triangle marker represents the single cutoff corresponding to delta. Delta-threshold refers to the baseline method that ranks CpG sites by methylation difference between groups. DSS ranks sites using Wald statistics. MAGIC includes both Bayes factor test (bf) and Wald test (wald) statistics.

The MAGIC Bayes factor test consistently achieved the highest AUC value, indicating stronger detection performance. Under a particularly challenging setting (delta = 0.05, 5× coverage, 3 components; **Fig. 3A**), all methods exhibited reduced sensitivity due to the weak signal. Nevertheless, MAGIC still reached an AUC of 0.632, highlighting its strong detection power even under low coverage, small effect size conditions. As expected, performance for all methods improved with increasing coverage and effect size (**Fig. 3B–C**), as higher sequencing depth provides more stable estimates of methylation proportions. When the number of mixture components increased to 5, the model captured more complex methylation patterns compared to the 3-component setting, but the overall detection performance remained largely consistent. The Wald test showed slightly improved performance, while the Bayes factor test exhibited a modest decline (**Fig. 3D**). This decline may be attributed to the reduced separability among mixture components as the number of components increases, leading to weaker evidence contrasts.

We further quantified overall model performance by summarizing AUC values across varying coverage levels (5×, 10×, and 20×), effect sizes (0.05 and 0.1), and number of components (3 and 5). As summarized in **Fig. 4**, MAGIC consistently achieved the highest AUC values. The performance advantage was most pronounced under challenging conditions and gradually converged with other methods as data quality improved.

**Fig. 4.**
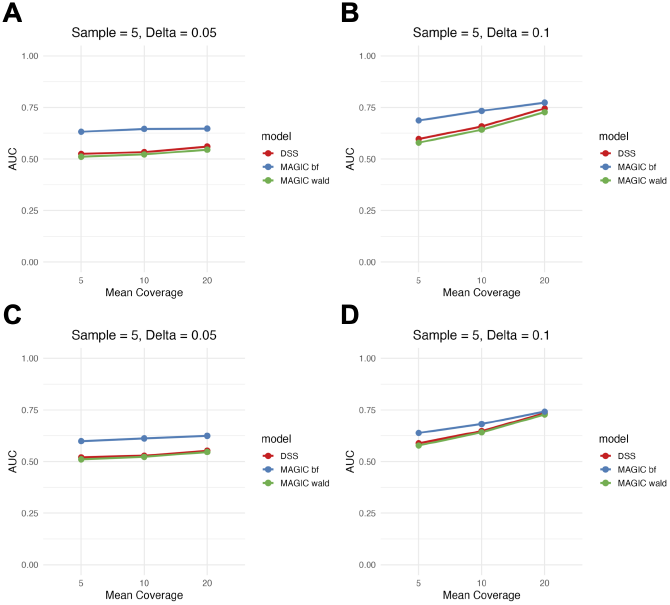
MAGIC Bayes factor test achieved highest AUC values under various simulations. (A) delta = 0.05, 3 components. (B) delta = 0.1, 3 components. (C) delta = 0.05, 5 components. (D) delta = 0.05, 5 components. All datasets included 5 samples per group.

DSS and MAGIC’s Wald test yielded comparable AUCs, whereas the MAGIC Bayes factor test showed a marked advantage under low-effect conditions (delta = 0.05; **Fig. 4A, C**), outperforming all other methods by more than 0.1 in AUC across coverage levels. Importantly, MAGIC’s advantage remained substantial even at 20× coverage, indicating that context-specific shrinkage provides benefits beyond simply compensating for low read counts. At a higher effect size (delta = 0.1) with 3 components, all methods showed improved absolute performance, but the relative ranking remained consistent (**Fig. 4B**). The performance tended to converge at 20× coverage, where all methods approached the highest AUC value at around 0.75.

Varying the number of mixture components exerted minimal influence on detection performance (**Fig. 4C–D**). For the MAGIC Wald test and DSS, performance across coverage levels was nearly identical between 3 component and 5 component datasets. This stability can be attributed to the fact that the Wald test evaluates differential methylation by comparing point estimates and their standard errors. In contrast, the MAGIC Bayes factor test showed slightly more sensitivity to the number of components, with modest performance degradation when increasing from 3 to 5 components. This likely reflects the increased parameter estimation uncertainty and the challenges in component assignment in the Bayesian model as data complexity rises. Nevertheless, even with 5 components, MAGIC Bayes factor maintained the highest performance among all competing methods across the conditions, indicating that the benefits of context-specific shrinkage outweigh the costs of increased complexity within reasonable ranges.

These comprehensive simulation results demonstrate that MAGIC provides consistent and substantial improvements in differential methylation detection across a wide range of realistic experimental scenarios, with the Bayes factor test offering particular advantages under challenging conditions of low coverage, small effect sizes, and limited replication.

## 4 Discussion

In this study, we developed MAGIC, a Bayesian mixture model designed to improve differential methylation detection under low coverage and limited replications for WGBS data. Our simulation results demonstrated that the MAGIC Wald test achieved well-calibrated p-values under null conditions. Notably, the MAGIC Bayes factor test consistently outperformed existing approaches, such as DSS, and baseline methods. These findings collectively highlight MAGIC’s strong detection power and well-controlled false positive rate across a wide range of sequencing depths and effect sizes, addressing a critical methodological gap in WGBS data analysis.

DNA methylation studies in WGBS data are often constrained by the dual challenges of low sequencing depth and small sample sizes, which together substantially reduce the statistical power to detect subtle but biologically meaningful methylation differences (Ziller *et al*. 2015; Feng *et al*. 2014). This power deficit leads to systematic underreporting of true differential methylation signals, which is particularly problematic in epigenome-wide association studies (EWAS) where effect sizes typically range from 5-15% (Rakyan *et al*. 2011). Shrinkage-based methods, such as DSS, have made important progress in addressing this challenge by employing genome-wide dispersion estimation to stabilize inference (Dolzhenko and Smith, 2014). MAGIC extends this approach by additionally capturing methylation heterogeneity through context-specific shrinkage. In our comprehensive analysis, the MAGIC Bayes factor test demonstrated markedly higher AUC values compared to existing methods under low-coverage conditions. This can be attributed to MAGIC’s context-specific shrinkage strategy, which adaptively estimates dispersion parameters within components rather than applying global shrinkage. These results indicate that adaptive mixture models can serve as a powerful strategy to recover statistical power in WGBS datasets.

We specifically compared MAGIC against DSS and a baseline method for two reasons. First, previous comprehensive benchmarks have demonstrated that DSS performs well with limited biological replicates (Piao *et al*. 2021). Second, DSS established the foundation of combining beta-binomial modeling with dispersion shrinkage, making it the natural benchmark for evaluating how MAGIC’s component-specific shrinkage extends this framework. The baseline method provides a non-statistical reference point for assessing the value of statistical modeling.

Traditional differential methylation analyses typically rely on fixed significance thresholds (e.g., FDR < 0.05), which become increasingly problematic with sparse data (Robinson *et al*. 2014; Efron *et al*. 2012). Under these conditions, a large proportion of true positives remain undetected because the statistical evidence is diluted by read-level variability and small sample sizes. Our ROC plots explicitly illustrate the inherent tradeoff between sensitivity and specificity. When stringent cutoffs are used to control false positive rates, sensitivity rapidly declines as coverage decreases, leading to under-detection of DMLs (Libertini *et al*. 2016). Conversely, relaxing the threshold increases detection power but also necessarily inflates false positive rates, complicating biological interpretation (Storey *et al*. 2003). This sensitivity-specificity tradeoff represents a fundamental challenge extending across genomic studies, where low counts and limited replication reduce the ability to distinguish true signals from sampling variability (Schurch *et al*. 2016; Robinson and Oshlack, 2010). In our simulation studies, DSS and baseline approaches suffered from this tradeoff more severely than MAGIC. Specifically, at 5× coverage with 5 samples per group and a 0.05 effect size, the MAGIC Bayes factor test demonstrated marked sensitivity-specificity balance compared to competing methods. When allowing equivalent false positive rates, the MAGIC Bayes factor test achieved approximately much higher true positive rates than DSS across varying thresholds. This advantage reflects MAGIC’s ability to more confidently distinguish true differential methylation from noise under challenging conditions, as evidenced by steeper ROC curves and higher AUC values.

Moreover, the inclusion of the Bayes factor test further enhances this advantage, providing an alternative for quantifying evidence that remains stable under sparse data where Wald tests can be unstable (Kass and Raftery, 1995). Wald tests provide well-calibrated *p* values for genome-wide FDR control, while Bayes factors offer interpretable evidence scales independent of sample size, facilitating cross-study comparisons and prioritization of candidates for experimental validation (Goodman, 1999). Beyond improving detection power, MAGIC’s mixture framework provides interpretable biological insights through its inferred latent contexts. The characteristic parameters, including mean methylation, dispersion, and component weight, offer a data-driven decomposition of the methylation landscape that can be linked to functional genomic elements through enrichment analysis (Kuleshov *et al*. 2016). CpG sites assigned to distinct mixture components can also be analyzed using existing tools such as ChIPseeker (Yu *et al*., 2015) and Cistrome (Zheng *et al*. 2018) to identify regulatory effects. The soft component assignment further enables identification of CpG sites with ambiguous context memberships, potentially flagging regions undergoing epigenetic transitions or exhibiting allele-specific methylation patterns.

Despite these strengths, several limitations warrant consideration. MAGIC currently offers pair-wise differential analysis, limiting its applicability to multi-group studies. Computationally, the optimization procedure requires multiple random initializations to address local optima, and convergence quality depends on training strategies. Looking forward, natural extensions include multi-group comparisons, integration of continuous covariates through regression on mixture parameters, and application to single-cell bisulfite sequencing, enabling inference of cell-type-specific methylation states (Kapourani *et al*. 2021). Incorporating hierarchical structures to account for sample dependencies and integrating hidden Markov models to leverage spatial correlation would broaden MAGIC’s applicability to complex study designs and expand it to DMR detection (Sun *et al*. 2016).

Overall, MAGIC provides a novel method that unites statistical rigor with biological interpretability. By combining context-specific shrinkage, mixture-based modeling, and a dual testing strategy, it captures the heterogeneity inherent in methylation data without sacrificing robustness or scalability. The ability to classify variance structures offers an additional dimension for biological discovery and enhances interpretive depth. These strengths make MAGIC a versatile tool for differential methylation analysis in WGBS data. Future extensions that incorporate multi-group comparisons and DMR detection will further expand its scope to the analysis of complex epigenomic datasets.

## Supplementary Data

Supplementary data are available at bioRxiv.

## Acknowledgements

This project used computational and storage services associated with the Hoffman2 Cluster, operated by the UCLA Office of Advanced Research Computing’s Research Technology Group.

## Author’s Contributions

M.T. developed the method. J.H. performed data simulation and model evaluation. M.P. supervised the project. All authors read and approved the final manuscript for publication.

