## Supplemental Figure for "MAGIC: Methylation Analysis with Genomic Inferred Contexts"

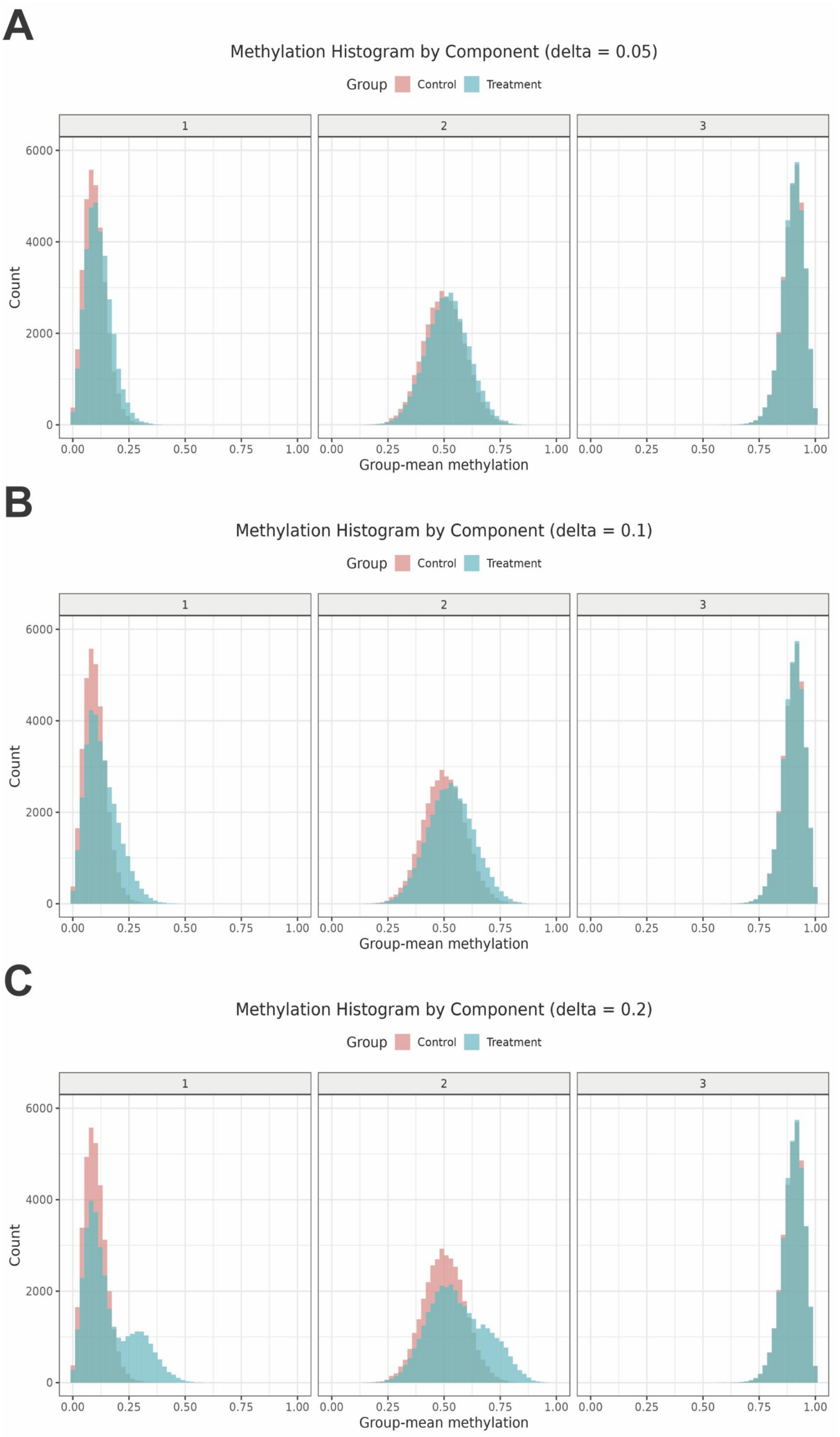


Supplementary Fig.1. Component-wise methylation distribution of simulated datasets with 3 components. (A) delta = 0.05. (B) delta = 0.1. (C) delta = 0.2. The three components follow beta distributions: Beta(10, 90), Beta(10, 10), and Beta(90, 10).

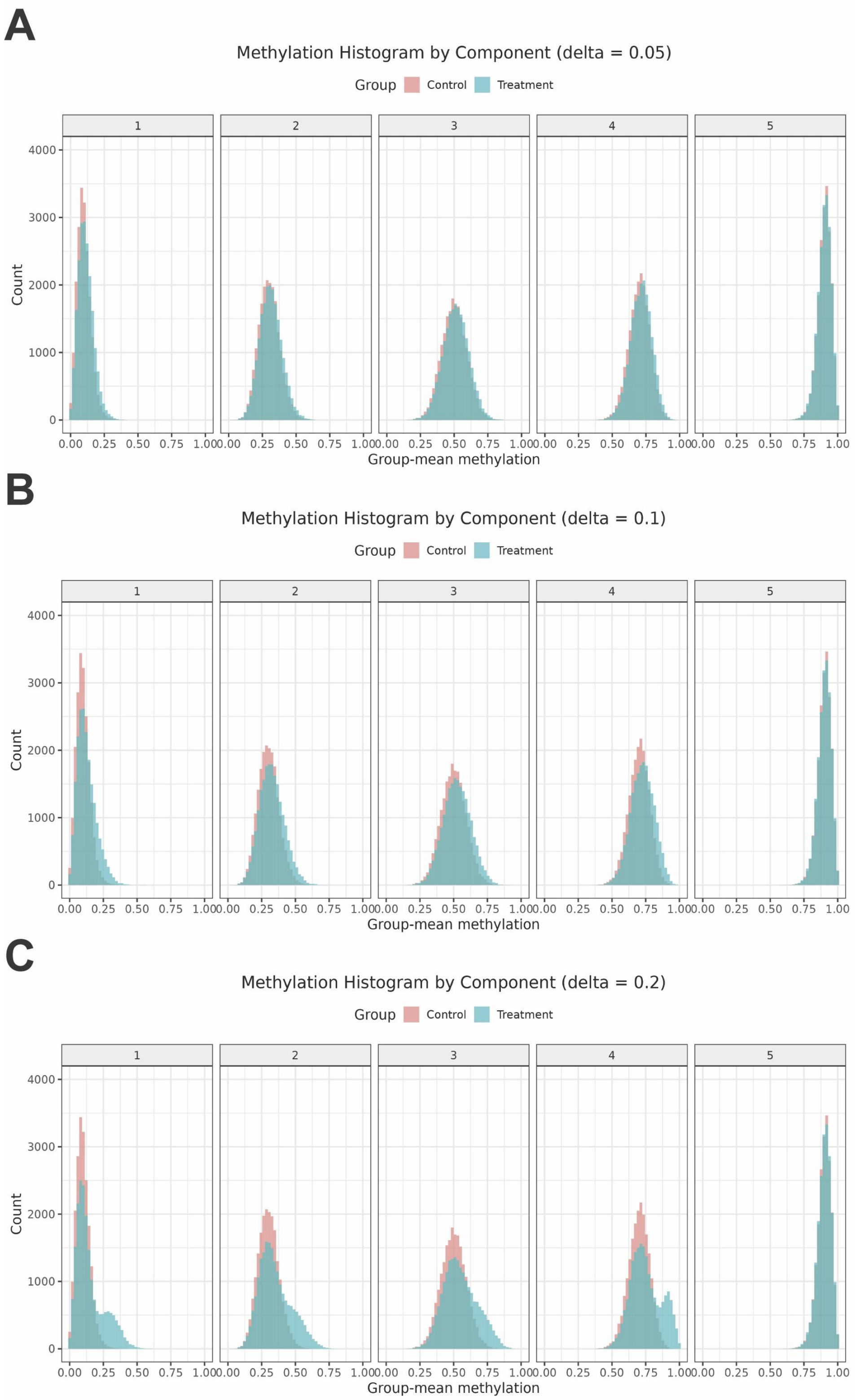


Supplementary Fig.2. Component-wise methylation distribution of simulated datasets with 5 components. (A) delta = 0.05. (B) delta = 0.1. (C) delta = 0.2. The five components follow beta distributions: Beta(10, 90), Beta(18, 42), Beta(10, 10), Beta(42, 18), and Beta(90, 10).

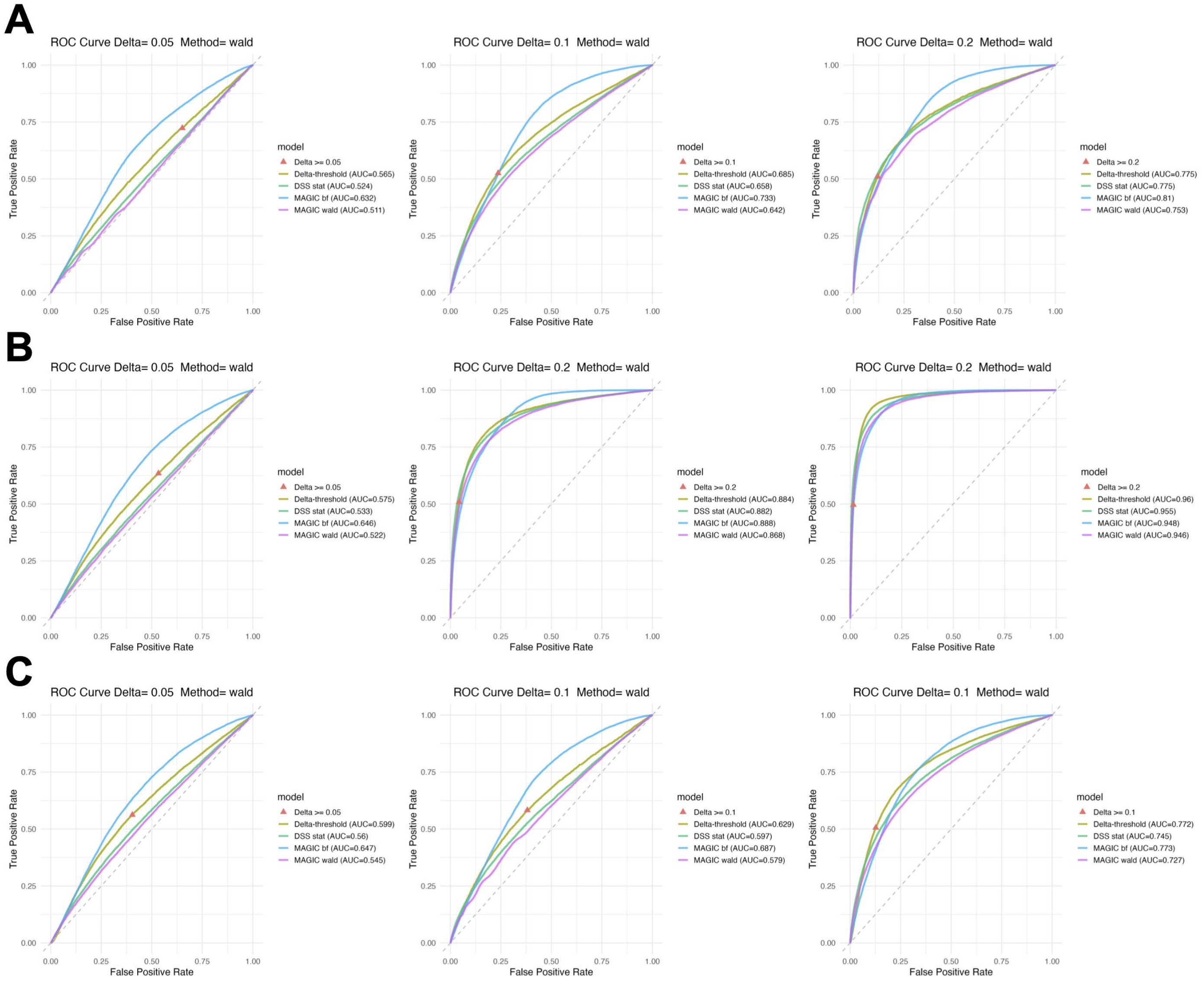


Supplementary Fig.3. Comparison of model performance on simulated datasets with 3 components. (A) 5x coverage. (B) 10x coverage. (C) 20x coverage. Each dataset includes 5 samples per group. The triangle marker represents the single cutoff corresponding to delta. Delta-threshold refers to the baseline method that ranks CpG sites by methylation difference between groups. DSS ranks sites using Wald statistics. MAGIC includes both Bayes factor test (bf) and Wald test (wald) statistics.

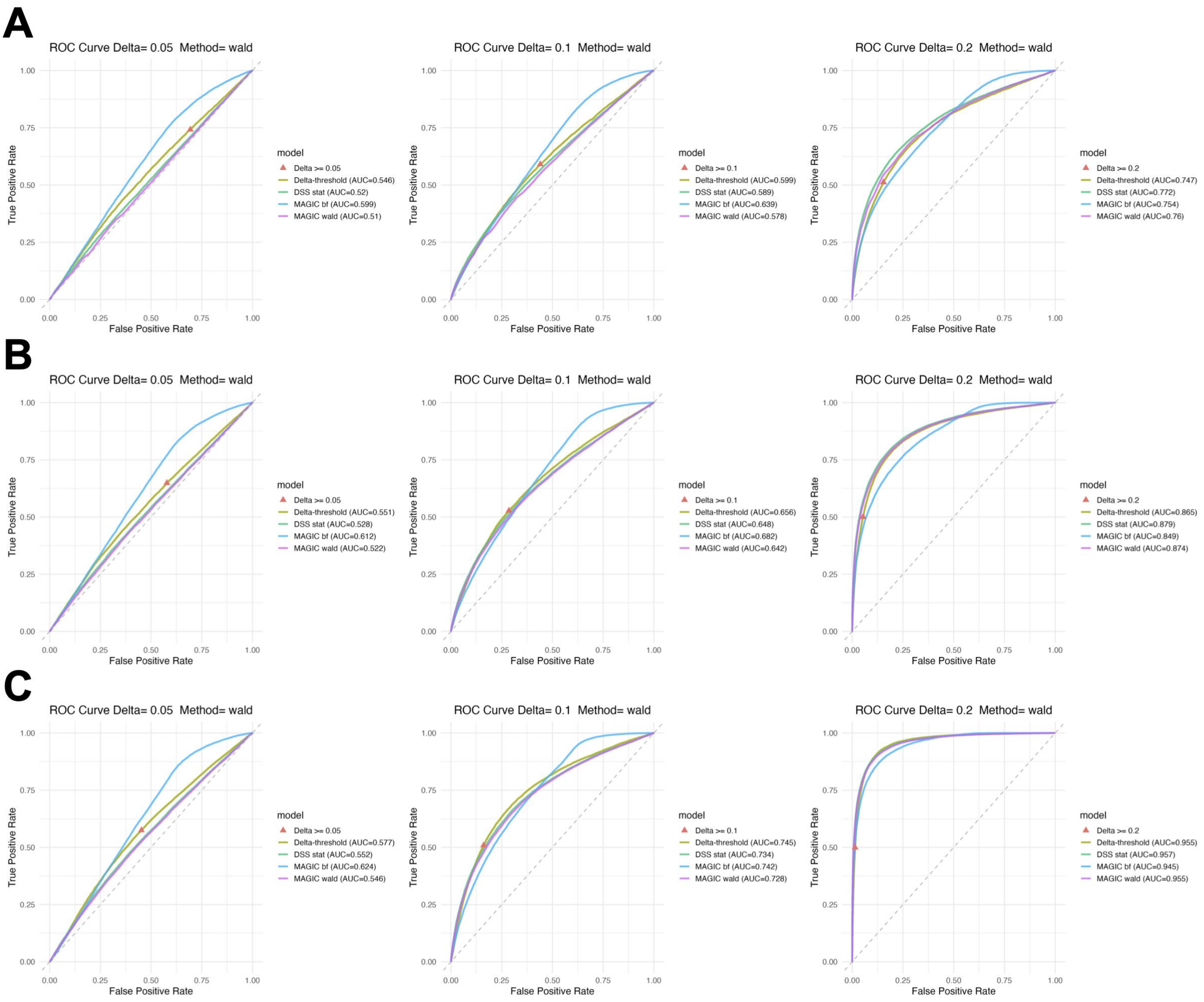


Supplementary Fig.4. Comparison of model performance on simulated datasets with 5 components. (A) 5x coverage. (B) 10x coverage. (C) 20x coverage. Each dataset includes 5 samples per group. The triangle marker represents the single cutoff corresponding to delta. Delta-threshold refers to the baseline method that ranks CpG sites by methylation difference between groups. DSS ranks sites using Wald statistics. MAGIC includes both Bayes factor test (bf) and Wald test (wald) statistics.
